# Host genetic variation in feeding rate mediates a fecundity cost of parasite resistance in a *Daphnia*-parasite system

**DOI:** 10.1101/2022.11.29.518345

**Authors:** Stuart K.J.R. Auld, Kätlin Raidma

**Author notes:** Corresponding author, **Correspondence address:** Room 3A149, Cottrell Building, University of Stirling, Stirling, FK9 4LA, UK. **Email:**.

## Abstract

Organisms face numerous challenges over their lifetimes, including from competitors and parasites, and experience selection to maximise their fitness in the face of these various pressures. However, selection can rarely maximise individual ability to cope with all challenges, and trade-offs therefore emerge. One such trade-off is the cost of resisting parasitic infection, whereby hosts that have a high intrinsic capacity to resist parasitic infection have comparatively low fitness in the absence of the parasite, and spatio-temporal variation in the relative strength of parasite- and non parasite-mediated selection is thought to maintain diversity in host resistance. Here, we test for, and find, a simple cost of resistance in the freshwater host *Daphnia magna* and its sterilising bacterial parasite, *Pasteuria ramosa* that is shaped by ecology as opposed to immunity. We uncovered significant genetic variation in *Daphnia* feeding rate, and show that rapid-feeding *Daphnia* genotypes have high fecundity in the absence of the parasite, but are more likely to go on to suffer sterilising infection when exposed to the parasite. This feeding rate-mediated cost of resistance can explain the persistence of parasite-susceptible genotypes. Further, we found evidence of infection induced anorexia in *Pasteuria*-infected hosts. It follows that reduced feeding in infected hosts means that high parasite prevalence could result in greater host food availability; this could reduce intra-specific competition and mask the cost of resistance in nature.

**Summary statement:** Costs of *Daphnia* immunity to a sterilising bacterial parasite are mediated by feeding ecology and not immunity, and infection-induced anorexia can further alter the relative strength of parasitism and host-host competition.

## INTRODUCTION

Infectious disease is a key challenge for nearly all organisms, and fitness is commonly shaped, at least in part, by an individual’s capacity to resist infection (Graham et al., 2011). Hosts have consequently evolved complex immune systems to identify infectious agents and coordinate immune effectors to destroy them (Frank, 2020). We know that fitness is only partly dependent on resistance because pathogen-susceptible genotypes persist in many host populations, and this strongly suggests that maintaining immune machinery comes at a cost to other fitness traits. Such costs could be energetic, or in the form of collateral damage of immune responses, or in some cases, increased susceptibility to an alternative pathogen species or even alternative genotype of the same pathogen species) (Graham et al., 2011; Schmid Hempel and Ebert, 2003; Sheldon and Verhulst, 1996). Although these constitutive costs of resistance are expected, there are many systems where they are either elusive, or only occur under very specific circumstances (see Labbé et al., 2010). For example, parasitoid-resistant *Drosophila melanogaster* have reduced larval competitive ability, but only when food is scarce (Fellowes et al., 1998; Kraaijeveld and Godfray, 1997), and whilst poultry selected for increased overall immunocompetence evolved reduced body mass, it was not the case that poultry selected for increased body mass suffered reduced immunocompetence (van der Most et al., 2011). What can explain the missing costs of resistance?

Resistance is not solely determined by the immune system, and experiments designed to target the effects of immunologically-driven resistance often eliminate variation in ecological conditions that make a larger contribution to anti-parasite defences (Lazzaro and Little, 2009). It follows that if we are understand the costs of resistance in more natural settings, we have to examine host genetic variation in non-immunological, and sometimes less obvious anti-infection mechanisms (Auld et al., 2013). For example, feeding rate is an obvious fitness trait that is also associated with exposure to environmentally-transmitted parasites. It has long been known that organisms that have high feeding rates will be successful inter- and intra-specific competitors and enjoy relatively high fitness due to their ability to acquire more resources (Burnet et al., 1977; Galimany et al., 2017). However, in environments where environmentally-transmitted parasites are either common or virulent, rapid feeders could pay a fitness cost in that they are more likely to encounter infectious parasite transmission stages and suffer the fitness implications of disease (Auld et al., 2013). Theory tells us that this simple trade-off tell us about the relative strength of competition and parasitism as ecological forces of selection, and could potentially explain the persistence of parasite-susceptible genotypes (Hall et al., 2010; Walsman et al., 2022); indeed, when competition is particularly strong, rapid feeding and higher susceptibility can be favoured by selection even when parasites are prevalent.

In many host-parasite systems, matters are further complicated by the fact that infected hosts suffer altered feeding rates. Infection induced anorexia is common and in some cases adaptive for the host (see (Ayres and Schneider, 2009). and references within). Moreover, in systems where infection induced anorexia occurs, an increase in parasite prevalence can result in greater food availability for the uninfected fraction of the host population, thus reducing intra-specific competition and potentially mitigating any cost of resistance (Walsman et al., 2022).

In this study, we used the crustacean *Daphnia magna* and its environmentally-transmitted sterilising bacterial parasite, *Pasteuria ramosa*, to test the following hypotheses: (1) genetic variation in *Daphnia* feeding rate is positively associated with in the absence of infection; (2) genetic variation in *Daphnia* feeding rate is also negatively associated with likelihood of suffering infection by *Pasteuria;* and (3) that feeding-rate is reduced in *Pasteuria-infected Daphnia.*

## MATERIALS AND METHODS

### Study system

The host, *Daphnia magna* is a common freshwater crustacean that inhabits shallow ponds throughout Europe. *Daphnia* are filter-feeders that graze on algae, but they also frequently ingest spores of horizontally-transmitted parasites, such as the bacterium *Pasteuria ramosa* (Auld, 2014; Ebert, 2008). After ingestion, *Pasteuria* spores bind to and then penetrate the *Daphnia* oesophagus (Duneau et al., 2011); after sporulation, the parasite undergoes a 10-20 day developmental process, and many millions of transmission spores are then released after host death (Ebert et al., 1996). The process of parasite within-host growth and development is very costly to the host, and infection commonly results in complete castration (Cressler et al., 2014; Ebert et al., 2004).

We conducted a suite of experiments to test the relationships between host feeding rates, host fecundity in the absence of infection and susceptibility to infection across 16 *Daphnia* genotypes (=clones). Host genotype lines were established from single individuals taken from 16 different experimental ponds at the University of Stirling (the Stirling Outdoor Disease Experiment; see Auld and Brand, 2017). These ponds were all initiated with the same suite of 12 *Daphnia* genotypes in 2015 (originally from Kaimes Farm, Leitholm, Scottish Borders (2°20’43’W, 55°42?5”N)), but the multiple rounds of sexual reproduction and thus genetic recombination means we are certain that *Daphnia* harvested from differing ponds are genetically distinct. The *Pasteuria* population we used comprised of a diverse mix of multiple isolates from the same original population as the *Daphnia.*

We maintained three replicate maternal lines for each of our *Daphnia* genotypes (a total of 48 maternal lines). Each replicate line was established with eight neonate (<24h old) *Daphnia* in jars containing 200mL of artificial *Daphnia* medium (ADaM: Klüttgen et al., 1994, modified using 5% of the recommended SeO2 concentration; Ebert et al., 1998). Each replicate line was fed 8 ABS of *Chlorella vulgaris* algal cells per day (where ABS is the optical absorbance of 650nm white light by the *Chlorella* culture). *Daphnia* medium was changed twice per week, or after the *Daphnia* produced offspring (the offspring were discarded). Maternal lines were maintained for three generations to minimise any variation due to maternal effects; the second and third generations were established using 2^nd^-3^rd^ clutch offspring. Experimental replicates were established using 2^nd^-4^th^ clutch offspring from each maternal line.

### Experimental 1: Genetic variation in feeding rate and fecundity

From each replicate maternal line, neonate (<24h old) offspring were harvested and allocated to experimental jars on day zero. All experimental jars contained 8 *Daphnia* in 200mL artificial *Daphnia* media (identical to the maternal lines), and were fed 8 ABS *Chlorella* algal cells per day and had media refreshed on day three.

On day six, a feeding rate assay was conducted. Media was refreshed and each experimental jar was fed 8 ABS *Chlorella* algal cells; the algae and media was thoroughly mixed with a plastic coffee stirrer (one per jar). An additional eight jars containing no *Daphnia* were also setup (and were fed the same algae and stirred also). A 1.5mL sample of the media was then taken and the optical absorbance was determined from each jar immediately after stirring (0 minutes), and then at 30, 60 and 120 minutes. Thus, we quantified algal densities over time in media *Daphnia*, and in media with no *Daphnia.* Immediately after the assay was completed, media was once again refreshed and *Daphnia* were fed 8 ABS *Chlorella* cells per day.

On days 8, 10, 12, and 16, the number of surviving adults and number of offspring were recorded in each jar; the offspring were then discarded, media refreshed, and the jars were fed 8 ABS per day.

### Experiment 2: Infection and feeding rate across genotypes

Experimental protocol was similar to that of Experiment 1. From each replicate maternal line, neonate (<24h old) offspring were harvested and allocated to experimental jars on day zero. All experimental jars contained 8 *Daphnia* in 200mL artificial *Daphnia* media (identical to the maternal lines). All jars received a dose of 1 x 10^5^ (50μL) *Pasteuria* transmission spores (comprising homogenized, previously infected *Daphnia* diluted in ddH_2_O). All replicate jars were fed 2 ABS *Chlorella* algal cells per day for a three-day (72h) exposure period. After the exposure period (day four), replicates were changed into new jars with fresh media and fed 8 ABS *Chlorella* algal cells per day. On day six, a feeding rate assay was conducted in a similar manner to that described for Experiment 1, except that the optical absorbance of media was determined at two time points: 0 and 120 minutes.

On days 8, 10, 12, and 16, media was refreshed and offspring were then discarded from each jar; the jars were then fed 8 ABS per day. On day 20, the number of *Pasteuria*-infected *Daphnia* within each jar was recorded (infected hosts are larger than their uninfected counterparts, red brown in colour and have an empty brood chamber due to parasite-induced castration). We then conducted a second feeding assay. All animals were placed into fresh jars and fed 8 ABS *Chlorella* algal cells following the protocol described previously and the optical absorbance of the media was determined at 0 and 120 minutes. Again, an additional eight jars containing no *Daphnia* were also setup (and were fed the same algae and stirred also) as controls.

### Analysis

Experiment 1 data were analysed as follows. We first tested whether algal density declined over time in the *no-Daphnia* (control) jars. This was done using a simple linear model fitted to control jar absorbance data, where sample time was fitted as a covariate. We then confirmed that the initial algal absorbance (*I.e.*, at time=0h) was the same across all genotypes and controls. This was done by fitting a linear model to data from both control and *Daphnia* samples, with genotype (genotype ID, or *no-Daphnia*) fitted as a fixed factor.

Next, we tested for genotypic variation in *Daphnia* feeding over time (excluding the no *Daphnia* control treatment). We did this by subtracting the absorbance at t=60 and t=120 from absorbance at t=0 for each jar; we then divided by 8 (the number of *Daphnia* per jar) to convert the value from being loss of algae in the media to being the cumulative absorbance of algae consumed per *Daphnia.* Next, we fitted a linear model to these feeding data, including genotype ID, sample time and their interaction as fixed effects.

Experiment 2 data were analysed in a similar manner to the data from experiment 1. For both the Day 6 and Day 20 feeding rate assays, we confirmed that the initial algal absorbance (*i.e.*, at time=0h) was the same across all genotypes and controls, and was unaffected by future proportion of hosts that suffered infection. This was done to confirm the assay methods were appropriate.

Using the Day 6 feeding rate assay data, we then tested the hypothesis that *Daphnia* early feeding rate was associated with an increased likelihood of them becoming infected; we also tested is susceptibility to infection varied across *Daphnia* genotypes. We did this by fitting a generalized linear model to the number of infected and healthy *Daphnia* on day 20, with *per capita* feeding rate and genotype ID fitted as fixed factors; a binomial error distribution was included in the model. Our final analysis was to test the hypothesis that infection status had an impact on feeding rate. This was done using a simple linear model fitted to the feeding rate data, where proportion of infected hosts was fitted as a covariate and genotype ID a fixed factor.

## RESULTS

### Experiment 1: Genetic variation in feeding rate and fecundity

We found no significant decline in the *C. vulgaris* absorbance over time in control (no-*Daphnia*) jars (linear model (LM): intercept = 1.063x 10^-3^, slope = 4.137x 10^-5^, *F*_1,22_ = 2.45, *P* = 0.13; Fig. S1A). There was also no significant variation in initial (t=0) absorbance among control (*i.e.*, no *Daphnia*) and all 16 genotype treatments (LM: *F*_15,40_ = 0.49, *P* = 0.93; Fig. S1B). We found that *Daphnia* consumed a mean of 3.55 x 10^-3^ ± 0.21 x 10^-3^ (standard error) ABS *C. vulgaris* cells *per capita* over 120 minutes. Cumulative consumption of food increased over time (effect of time, LM: *F*_1,112_ = 259.07, *P* < 0.0001, 55% variation explained) and there was genetic variation in food consumed (effect of genotype ID, LM: *F*_15,112_ = 4.27, *P* < 0.0001, 14% of variation explained); cumulative consumption also varied according to genotype (effect of a time x genotype ID interaction, LM *F*_15,112_ = 2.49, *P* = 0.003, 8% of variation explained; Fig. 1A), thus demonstrating limited but nonetheless significant genetic variation in the profile of *Daphnia per capita* feeding rate. We also uncovered significant genetic variation for *Daphnia* early reproductive rate (effect of genotype ID, LM: *F*_15,224_ = 6.84, *P* < 0.0001, Fig. 1B). Finally, we found a significant positive relationship between host feeding rate (the slope of *Daphnia per capita* cumulative feeding over time) and *Daphnia* early reproductive rate (LM: *F*_1,14_ = 12.97, *P* = 0.003; Fig. 1B)

**Figure 1.**
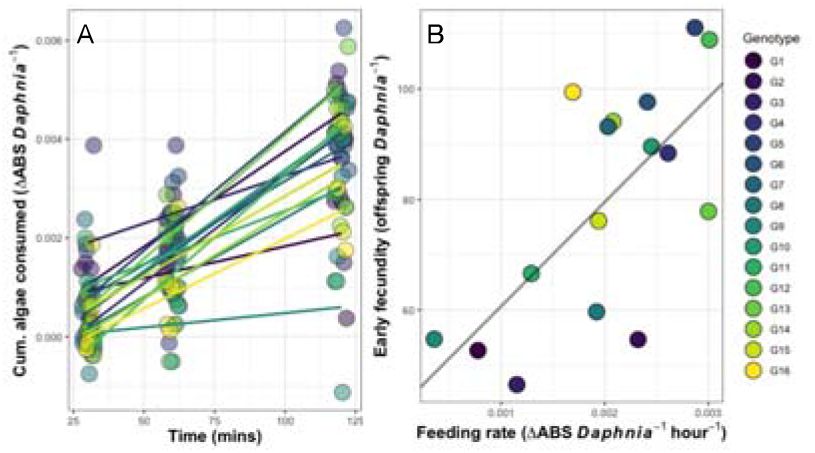
A Genotypic variation in Daphnia Cumulative consumption of C. vulgaris algae over 120 minutes. B Relationship between Daphnia feeding rate and early fecundity across. Line (A) and point (B) colours denote Daphnia genotype ID.

### Experiment 2: Infection is a consequence and cause of variation in feeding rates

On day 6, there was no significant variation in initial (t=0) absorbance among control (*i.e.*, no *Daphnia*) and all 16 genotype treatments (LM: *F*_16,38_ = 0.39, *P* = 0.98; Fig. S2A), nor was there variation in initial absorbance according to the proportion of *Daphnia* in each jar that went on to suffer infection (LM: *F*_1,38_ = 0.23, *P* = 0.63; Fig. S2B). When we analysed algal consumption data from jars that contained *Daphnia*, we found that average *per capita* feeding rate was positively associated with the proportion of *Daphnia* that went on to suffer infection (effect of proportion infected, GLM: χ^2^_1_= 18.94, *P* < 0.0001; Fig. 2A) and varied according to *Daphnia* genotype /effect of genotype ID GiM: χ2 = 3106 *P* = 0007; Fig 2B).

**Figure 2.**
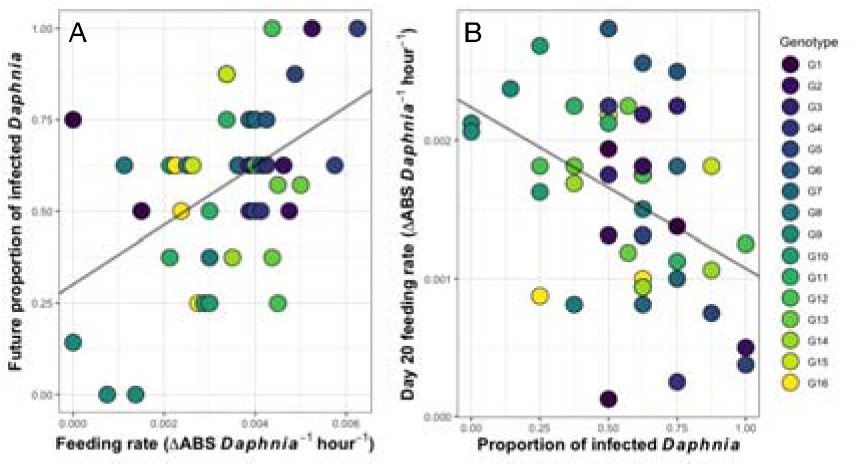
**A** Relationship between Day 6 *Daphnia* feeding rate and future proportion of hosts infected. **B** Relationship between proportion of hosts infected and Day 20 *Daphnia* feeding rate Daphnia feeding rate and early fecundity across. Point colours denote *Daphnia* genotype ID.

On day 20, there was no significant variation in initial (t=0) absorbance among control (*i.e.*, no *Daphnia*) and all 16 genotype treatments (LM: *F*_16,38_ = 1.11, *P* = 0.38; Fig. S2C), nor was there variation in initial absorbance according to the proportion of *Daphnia* in each jar that went on to suffer infection (LM: *F*_1,38_ = 0.05, *P* = 0.82; Fig. S2D). We did find that infection status had a strong impact on feeding rate in parasite-exposed replicates: mean *per capita Daphnia* feeding rate decreased with an increasing proportion of infected animals (effect of proportion infected *Daphnia*, LM: *F*_1,31_ = 11.28, *P* = 0.002; Fig. 2B), though there was now no significant variation in *Daphnia per capita* feeding rate according to genotype ID (effect of genotype ID, LM: *F*_15,31_ = 1.66, *P* = 0.12).

## DISCUSSION

Numerous studies have evaluated the costs of evolving and maintaining strong immune defences, and often either found none, or else found such costs under specific conditions (see Labbé et al., 2010 for a review). We tested for a simple non-immunological constitutive cost of resistance in the *Daphnia-Pasteuria* freshwater host-parasite system. Specifically, using two controlled laboratory experiments undertaken with sixteen genotypes of *Daphnia* and a mixed population of *Pasteuria* spores, we tested if host genetic variation in feeding rate mediated a fitness cost of resistance, and if there was evidence for reduced feeding in infected hosts, consistent with infection induced anorexia (Ayres and Schneider, 2009).

We found that algal densities declined over time in the presence of *Daphnia*, and that there was no significant decline in control replicates (where *Daphnia* were absent), thus demonstrating that *Daphnia* consume the algae. We uncovered significant genetic variation algal consumption, and that there was more limited (but nonetheless significant) genetic variation in feeding profiles over time: some genotypes slowed their consumption whilst others increased their feeding rate. Still, it is important to note that genetic variation in feeding rate profile accounted for a small proportion of the variation in the data compared to overall genotype variation in consumption, and this is supported by the limited changes in the rank order of genotypes over time. We therefore argue that 120 minute feeding rate is a useful measure to compare across *Daphnia* genotypes.

This variation in feeding rate was found to have important consequences for *Daphnia* fitness in the absence of infection. Not only did we uncover significant genetic variation for *Daphnia* early fecundity, we also found that early fecundity was strongly positively associated with feeding rate. Genotypes that rapidly filtered algae over a two hour period produced more than twice as many offspring than their slow feeding counterparts. All else being equal, rapid feeding genotypes would sequester the most resources, produce the most offspring and increase in relative frequency; rapid feeders would thus swiftly dominate populations on account of their intra-specific competitive ability.

However, results from our second experiment demonstrated that all else is not equal: we found that in the presence of the *Pasteuria* parasite, *Daphnia* genotypes with the greatest feeding rate were also most likely to go on to suffer infection. The most likely candidate explanation for this is that rapid feeding host genotypes have the greatest contact rate with *Pasteuria* transmission spores and would thus be exposed to a greater effective parasite dose rate. The cost of resistance is not a cost of immunity as is often expected, but a cost in terms of increased host exposure to parasite infectious stages. In the *Daphnia-Pasteuria* system, the probability of infection depends on the precise combination of host and parasite genotypes (Luijckx et al., 2011). Still, in natural environments *Daphnia* would encounter multiple *Pasteuria* genotypes, and experiencing a greater dose rate would therefore increase the likelihood the *Daphnia* encounter a *Pasteuria* spore that genotypically matches and causes infection (Ben Ami et al., 2008). Thus, in environments where *Pasteuria* transmission stages were abundant, the fitness advantage benefits with greater resource acquisition would be rapidly overshadowed with the massive fitness cost of suffering infection with a sterilising parasite; parasitism would come to outweigh competition as the dominant ecological force.

Finally, we found that replicate jars containing a greater proportion of *Daphnia* with established infections had a lower *per capita* feeding rate. This provides compelling evidence for *Pasteuria* induced anorexia in *Daphnia.* In many ways, this is surprising. Previous work has demonstrated that both *Daphnia* and *Pasteuria* are resource limited, and that *Pasteuria* infection causes gigantism in its castrated hosts. It was argued that this gigantism was likely an adaptive parasite trait to sequester host resources for *Pasteuria* reproductive potential (Ebert et al., 2004). Put simply: *Pasteuria* steals future *Daphnia* offspring to fuel parasite spore production. However, infection induced anorexia could act counter to this. One explanation could be that *Daphnia* reduce their feeding rate upon infection in order to modulate their own life history in ways to mitigate the (massive) costs of infection. Infection induced anorexia could be the mark of the infected host shifting resource allocation away from survival/maintenance towards producing offspring more rapidly before castration is completed (termed fecundity compensation). Indeed, we observe fecundity compensation in multiple systems (Barribeau et al., 2010; Thornhill et al., 1986), including *Daphnia*-parasite systems (Chadwick and Little, 2005; Vale and Little, 2012). For *Daphnia-Pasteuria* interactions, fecundity compensation is viewed as a form of tolerance (as opposed to resistance) because it maintains host fitness without reducing the within-host parasite burden.

Infection induced anorexia could also have broader ecological consequences. The reduced food consumption in infected *Daphnia* means that increases in parasite prevalence could result in relaxed competition among host genotypes (including slow-feeding parasite resistant genotypes), thus masking the feeding cost of resistance. However, these hypothetical scenarios are, by their very nature, speculative and require rigorous testing under ecologically complex conditions. Similarly, we must also acknowledge that other factors affect *Daphnia* feeding rate: in the related *Daphnia dentifera* host, switching to a low quality food resulted in a reduced feeding rate, slower growth and reduced susceptibility to the *Metschnikowia bicuspidata* yeast parasite (Penczykowski et al., 2014). Nevertheless, it is clear that host feeding rate can be instrumental as both a cause and a consequence of parasitic infection.

## Acknowledgements

We thank NERC Iapetus2 for funding a Research Experience Placement to support KR.

## Author contributions

SKJRA conceived of the study; SKJRA and KR collected and analysed the data; SKJRA and KR wrote the manuscript. Both authors approved the final version of the manuscript.

## Data availability statement

All data will be archived on Dryad upon acceptance of the manuscript.

## SUPPLEMENTARY MATERIAL

**Figure S1.**
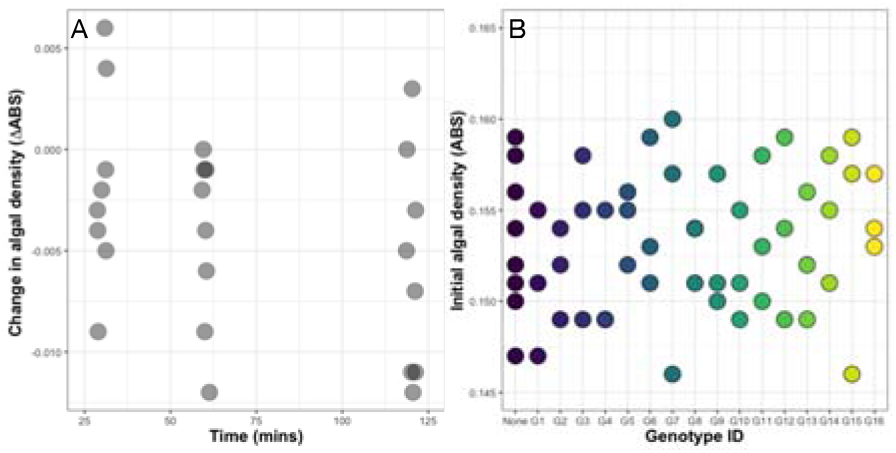
**A** No change in *C. vulgaris* algal absorbance over 120 minutes. **B** No difference in initial (t=0 mins) *C. vulgaris* algal absorbance across control (*Daphnia* = “None”) and 16 genotypes.

**Figure S2.**
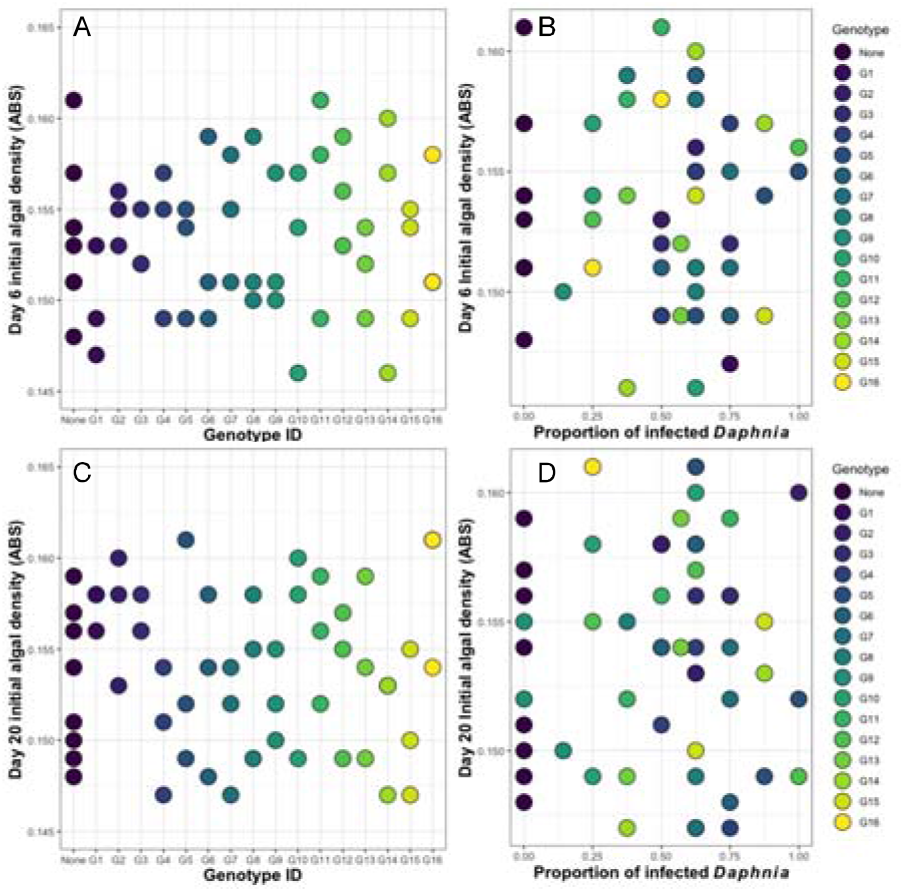
No variation in Day 6 initial *C. vulgaris* algal absorbance over 120 minutes t **A** across control (*Daphnia =* “None”) and 16 genotypes, and **B** between proportion of hosts that became infected. Also no variation in Day 20 initial *C. vulgaris* algal absorbance over 120 minutes **C** across control (*Daphnia* = “None”) and 16 genotypes, and **D** between proportion of hosts that became infected.

